# Ecological occurrence and plant regeneration of embryoid of the endangered and endemic plant *Dysosma versipellis* in China

**DOI:** 10.1101/471631

**Authors:** Xiao-Ming Tan, Ya-Qin Zhou, Pei-Bin Wu, Ying Wei, Shi-Lin Yang, Hong-Zhen Tang

## Abstract

In this study, the effective callus culture, somatic embryogenesis, and plant regeneration system of *Dysosma versipellis*, which is an endangered and endemic plant in China, were established under specific culture conditions. Using the *D*. *versipellis* leaves, petioles, and roots as explants, DPS software orthogonal design method and SPSS Duncan’s multiple range test were used to investigate their effects of *D*. *versipellis* on callus formation, embryoid induction, and plant regeneration by adding different phytohormones. Results showed that leaves and petioles were the most suitable materials in inducing callus. The effect of phytohormone on callus formation followed the order of 2,4-dichlorophenoxyacetic acid (2,4-D)>thidiazuron (TDZ)> kinetin>naphthylacetic acid (NAA)>2-ip. The best medium for callus formation was MS+2,4-D 1 mg/L+NAA 0.05 mg/L+TDZ 0.5 mg/L+2-ip 1 mg/L. The optimal medium to induce the formation of granular callus embryoid was MS+0.5 mg/L 6-BA+0.1 mg/L NAA, and the induction rate was 71.33%. The embryoid rooting and plant regeneration medium was MS+0.5 mg/L IBA+0.5 mg/L GA3. The optimal medium formula obtained in this study was suitable for the rapid induction of callus, embryoid, and plant regeneration of *D. versipellis* under in vitro culture conditions. Further study on the action mechanism, signal regulation mechanism, and artificial seed production of fungal elicitors affecting the accumulation of podophyllotoxin is important.

## Introduction

*Dysosma versipellis* (Hance) M. Cheng, which is an endangered medicinal plant in China, is a perennial herb of the family Oleaceae that grows under hillside forests, shrubs, shady and wet areas near the streams, bamboo forests, or under limestone evergreen forests at an altitude of 300–2400 m [1]. *D. versipellis* (Hance) M. Cheng is a commonly used traditional Chinese medicine that can clear away heat, detoxify the body, and dissipate phlegm and swelling [2]. This plant is also the natural source of the precursor for artificial synthesis of anticancer drugs, such as etoposide (VP-16) and teniposide (VM-26; i.e., podophyllotoxin) [3, 4]. At the same time, the leaves of *D. versipellis* are wide, the edges are 4–9 m shallow or deep, and the flowers are bright. Therefore, *D. versipellis* is a potted flower plant with considerable ornamental value.

Given the destruction of the habitat environment, low natural reproduction rate, and predatory exploitation of wild resources, the wild resources of *D. versipellis* are on the verge of exhaustion. *D. versipellis* has been included in the first list of rare and endangered plants in the country as a third-class protected plant [5]. With the continuous development of new functions of podophyllotoxin and the increasing cancer incidence rate in humans [6], the contradiction between the supply and demand of *D. versipellis* is prominent, and carrying out the effective rapid propagation research on *D. versipellis* is important to solve the contradiction between the further development and effective protection of the source materials of podophyllotoxin [7, 8]. Researchers have conducted studies on the induction, differentiation, and proliferation of the calla of *D. versipellis*. However, study on the somatic embryogenesis and plant regeneration of *D. versipellis* has not been conducted to date. The formation of regenerated plants by somatic embryogenesis in an in vitro culture is an extremely common phenomenon [9-11]. The successful induction of plant somatic embryoids can provide a good system to study the metabolic mechanism of active components, differentiation and development of the plant cells, omnipotent expression, variety improvement, and mutant screening of endangered *D. versipellis*, which is important both in theory and application. Therefore, the orthogonal design method was used in the present study to explore the effects of different plant hormones on the callus induction and embryoid formation of *D. versipellis* with its leaves and petioles. This study aims to provide a database for the development of an efficient, stable, high-frequency, and rapid propagation program for *D. versipellis* through somatic embryogenesis.

## Materials and methods

### Plant materials

The leaves, petioles, and roots used in the experiment were all obtained from the tissue culture seedlings cultivated in our laboratory. The wild plants, which were identified as *D. versipellis* (*D. versipellis* [Hance] M. Cheng ex Ying) by Dr. Tan Xiaoming (Fig. 1a, b), were collected from Yongfu, Guilin, Guangxi Zhuang Autonomous Region, China (109°36’E, 24°37’N).

**Fig. 1.**
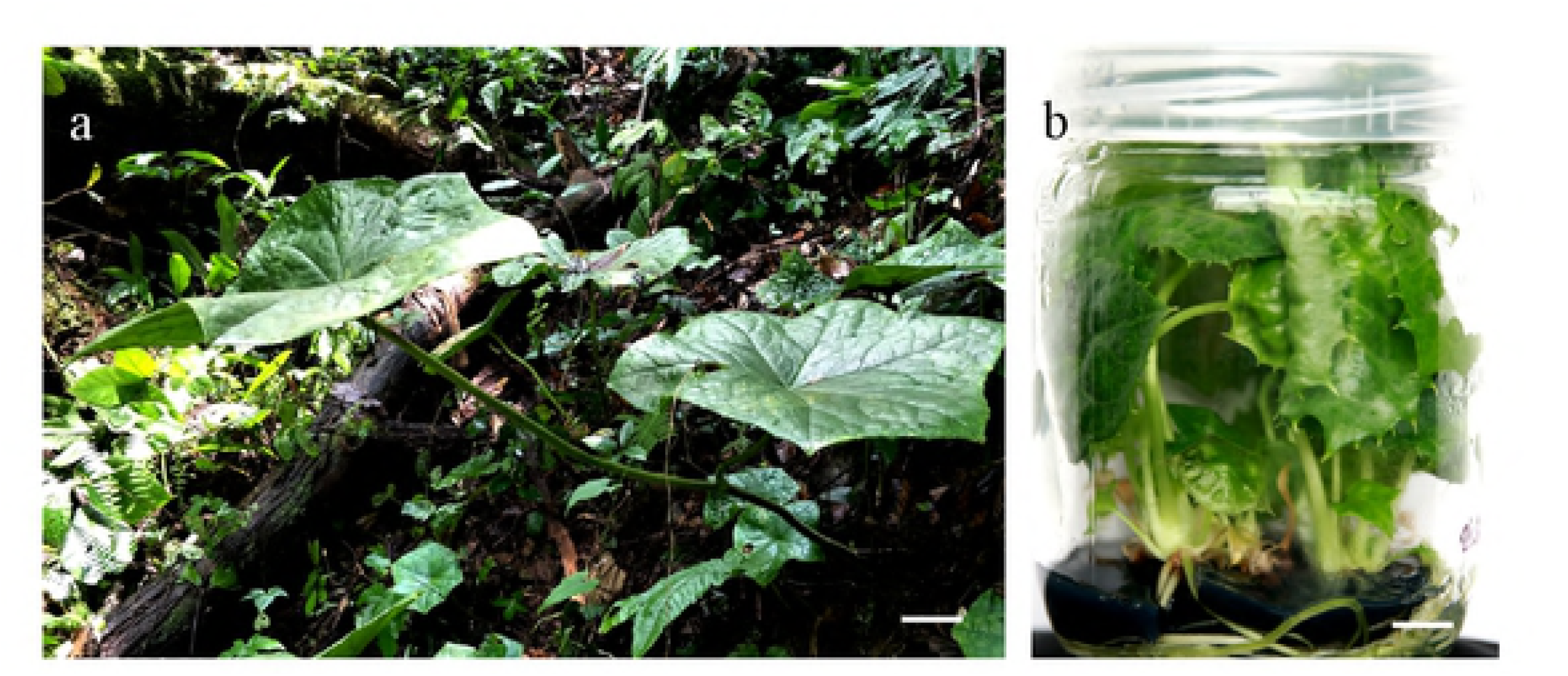
Plants and habitats of wild *Dysosma versipellis* (a;bar=10 cm) and their tissue culture seedings (a;bar=5 mm)

### Culture conditions

MS was used as the basic medium. Then, 30 g/L sucrose, 7 g/L agar, and 1 g/L activated carbon were added. The pH was adjusted to 5.8. Plant growing substances were added according to the requirements of the experiment. The culture temperature was 24 °C±2 °C. Callus induction and embryoid formation required dark culture, and root induction and plant regeneration required 10–12 h/day light culture with the light intensity of 30 μmol·m^−2^·s^−1^.

### Explant treatment

The vigorously grown *D. versipellis* seedlings were selected (Fig. 1b). The leaves, petioles, and roots were cut into small pieces (segments) of approximately 5 mm on the sterile and clean workbench and then inserted into the MS medium containing the plant growing materials.

### Leaf callus induction

The orthogonal test of L_16_(4^5^) was designed using the orthogonal design function of the DPS software. Taking *D. versipellis* leaves as explants, the effects of five growth substances 2,4-dichlorophenoxyacetic acid (2,4-D), thidiazuron (TDZ), naphthylacetic acid (NAA), kinetin (KT), and 2-ip on callus induction were explored. Four bottles of each treatment were prepared, and 4 to 5 pieces per bottle were inoculated to conduct regular observation. After 30 days, the visual statistical analysis of the induction rate, browning rate, and growth status was carried out.

### Somatic embryo induction

According to the subcultured callus, the vigorously growing granular and yellowish calli were selected and inoculated with four MS mediums, that is, 2,4-D 1 mg/L+KT 0.1 mg/L, 2,4-D 0.5 mg/L, 2,4-D 0.5 mg/L+NAA 0.1 mg/L, and 6-BA 0.5 mg/L+NAA 0.1 mg/L, to investigate the effects of 2,4-D, 6-BA, and NAA on embryoid induction. Four bottles of each treatment were prepared, 4–5 pieces per bottle were inoculated for regular observation. The statistical analysis of the induction rate was performed for 30 days for 8 times.

### Root induction and plant regeneration

The normally developed *D. versipellis* embryos with cotyledons were transferred to MS+0.5 mg/L IBA+0.5 mg/L GA3 medium for the light/dark alternating culture (light culture for 10h/dark culture for 14 h) to induce root formation and plant regeneration.

### Statistical method

This study used DPS version 7.05 data processing system and SPSS version 19.0 statistical software to statistically analyze the data. First, the data of the callus induction rate were inputted into the data column of the orthogonal design L_16_(4^5^) table of DPS, and the orthogonal design statistical analysis was performed to obtain the range R value. The larger the R value was, the more evident the induction effect was. The incidence of embryoid was analyzed by variance analysis by conducting Duncan’s multiple range test on SPSS. The significance of the difference between the mean values of the induction rates of different treatment groups was detected, where P>0.05 indicated significant difference. P>0.01 showed extremely significant difference, and the data were expressed as mean induction rate±standard error.

## Results

### Selection of the suitable inducing callus explants

In this study, the dedifferentiation abilities of *D. versipellis* leaves, petioles, and roots were pretested using MS+TDZ 0.5 mg/L+NAA 0.1 mg/L. The results showed that the callus induction rates of leaves, petioles, and roots were 69.4%, 53.8% and 18.95%, respectively. The two calli were induced, and the first callus was differentiated from leaves, which was vigorously divided, loose, and fragile (Fig. 2-a). The long-term cultured browning callus will produce multiple trumpet-like embryoids (Fig. 2-b), which will further develop into complete plants with cotyledons and roots (Fig. 2-c). This result provides an idea for the induction of callus embryoids for the next step of the study. The second callus was differentiated from the stem segmentation, which was large and tightly packed (Fig. 2-c). During the dark culture, some calli differentiated into the germs (Fig. 2-d) and multiple white roots (Fig. 2-e). Root-induced callus exhibited severe browning. Therefore, the leaves or petioles were selected as the inducing material for callus for the next experiment.

### Effects of different hormones on the callus induction rate of the *D. versipellis* leaves

The callus induction rates of *D. versipellis* leaves under the different combinations of TDZ, NAA, 2,4-D, KT, and 2-ip were obtained via visual analysis by using orthogonal experiments designed by DPS software. After approximately 2 weeks of culture, the middle part of the leaves was bulged, the edge of the pieces was attached to the medium, and the incision was swollen, which gradually formed the callus. Table 1 shows that the callus induction rate of treatment 10 was 66.67%, the browning rate was only 11.11%, and the callus growth was also optimal. The induced callus, which is a good material to further induce *D. versipellis* embryoids, was mostly yellowish and granular. Although the induction rate of treatment 13 was high (70%), *D. versipellis* leaves easily turned brown in color at the later stage, with the browning rate of 20%, and the growth of callus was worse than that of treatment 10 (Fig. 2 f–j).

**Table 1.**
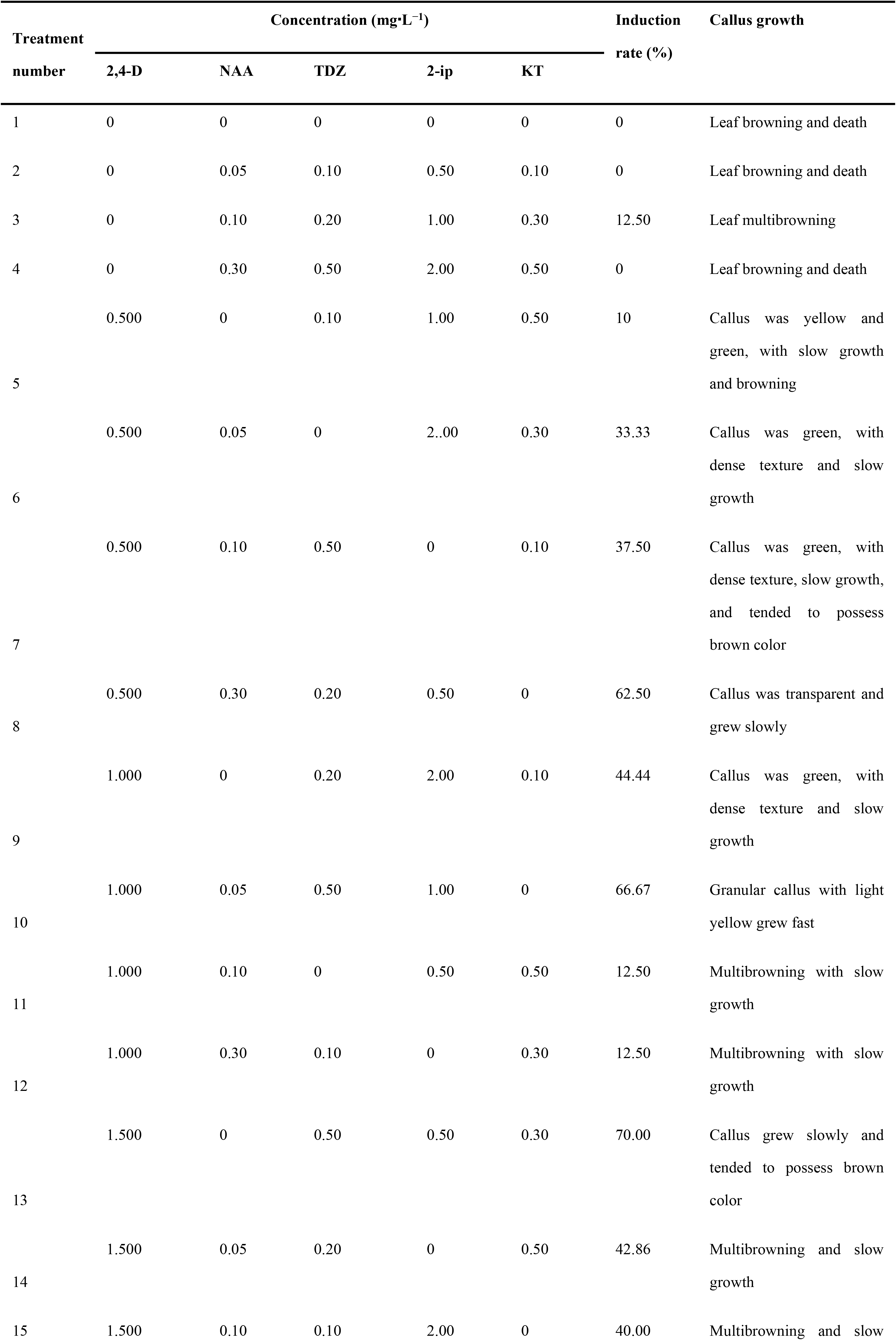
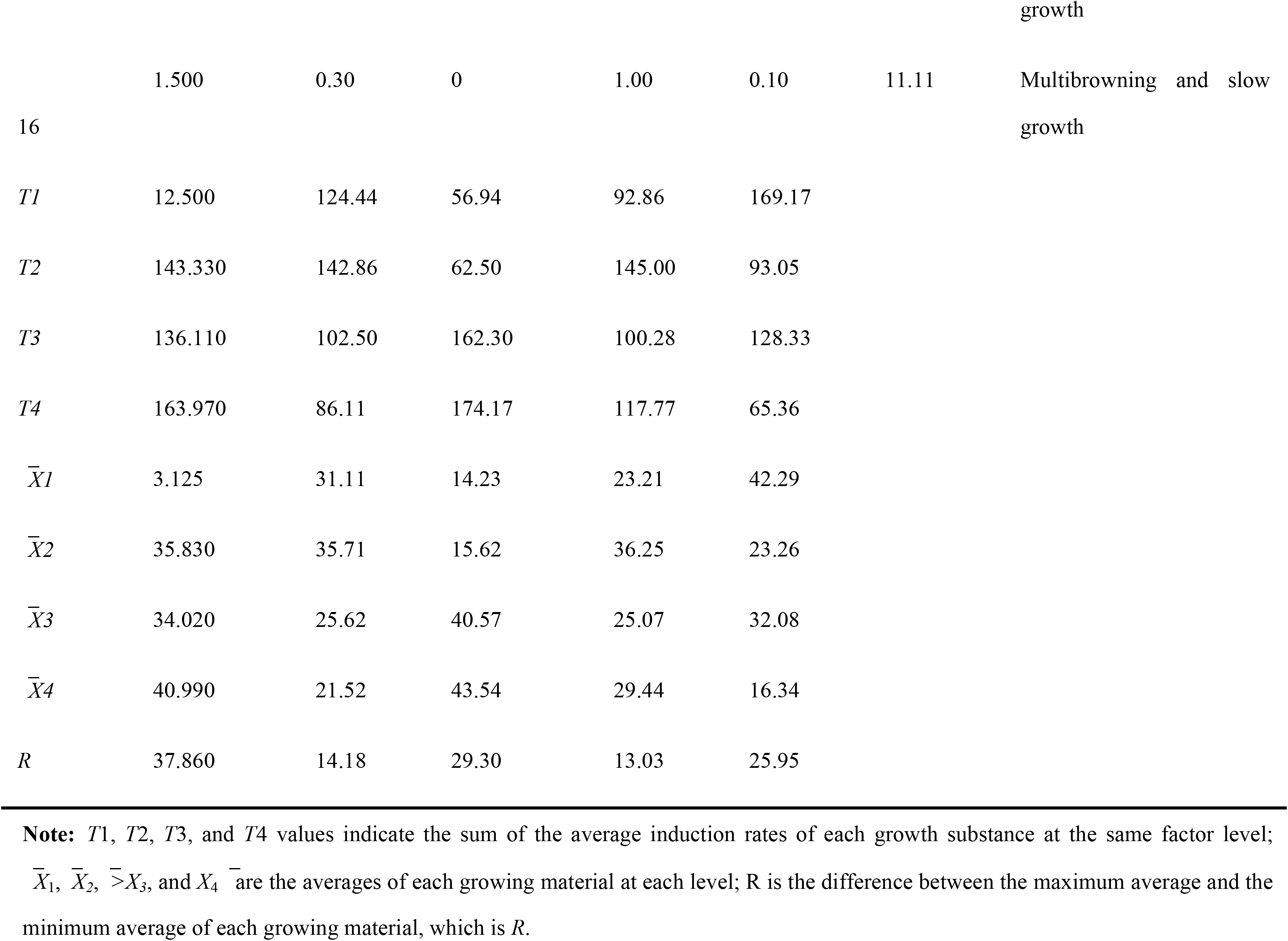
Orthogonal design of *Dysosma versipellis* leaf-induced callus and induction results

According to the statistical analysis method of DPS, the difference between the maximum and minimum means of callus induction rate of different growth hormones on *D. versipellis* was R; the larger the value of R was, the more evident the effect on the callus induction will be. Comparing the range R values of the five growing substances in the table and combining the growth status of callus showed that the effects of 2,4-D, NAA, TDZ, 2-ip, and KT on the induced callus in *D. versipellis* leaves followed the order of 2,4-D>TDZ>KT>NAA>2-ip. Among these hormones, 2,4-D and TDZ were the key factors affecting the callus formation of *D. versipellis* leaves, followed by KT, NAA, and 2-ip.

Therefore, a treatment combination for inducing the best growth material for the callus of *D. versipellis* leaves can be obtained, that is, the concentration of each growing substance when the total induction rate (T value) or the average number X value was the largest, as follows: MS+2,4-D 1 mg/L+ NAA 0.05 mg/L+ TDZ 0.5 mg/L+2-ip 1 mg/L.

### Effects of different hormone combinations on the occurrence of callus embryos in *D. versipellis*

After 8 generations of dark culture (1 subculture for 30 days), the *D. versipellis* callus can successfully induce spherical callus with granular surface on media 2, 3, and 4 (Fig. 2k) and gradually developed into different embryoid stages. On the basis of the whole development time, the occurrence of embryoid was unsynchronized. Despite the variation in sizes, they were mostly trumpet-like embryoids on the same callus (Fig. 2 l-n). These embryoids were loose and easily separated from the callus. Medium 1 showed insignificant embryoid formation. Using the Duncan’s multiple range test performed on the SPSS analysis software, the ANOVA of the average embryoid incidence in the callus of *D. versipellis* was carried out on different hormone combinations. The results showed that when the 2,4-D concentration was 0.5 mg/L, the average incidence rate of embryoid was 50.72% (Table 2). The results of the culture media 2 and 3 showed that when the 2,4-D concentration was 0.5 mg/L, the average incidence rate of embryoids was significantly higher than that of the medium without NAA by adding the medium with low NAA concentration (0.1 mg/L). The results of the culture media 3 and 4 showed that when the NAA concentration was 0.1 mg/L, the average incidence rate of embryoids under low 6-BA concentrations was significantly higher than that under low 2,4-D concentrations. By contrast, the combination of 6-BA and NAA with low concentration showed high activity on the callus formation of *D. versipellis*, and the most suitable subculture medium for callus embryoidogenesis was MS+0.5 mg/L 6-BA+0.1 mg/L NAA.

**Table 2.** Effects of different hormone combinations on the callus embryogenesis in *Dysosma versipellis*

| Treating number | Different hormone combinations and concentrations (mg/L) | Embryoid incidence rate (%) |
| --- | --- | --- |
| 1 | 2,4-D 1 mg/L+KT0.1 mg/L | 0 |
| 2 | 2,4-D 0.5 mg/L | 50.72±0.85<sup>Bc</sup> |
| 3 | 2,4-D 0.5 mg/L+NAA 0.1 mg/L | 52.33±1.52<sup>Bb</sup> |
| 4 | 6-BA 0.5 mg/L+NAA 0.1 mg/L | 70.70±1.54<sup>Aa</sup> |

**Note:** Different capital letters after the same number indicated that the difference was extremely significant ( $P<0.01$ ), and different lowercase letters indicated significant difference ( $P<0.05$ ).

MS+0.5 mg/L 6-BA+0.1 mg/L NAA can be used as a subculture medium for embryoids. If the conditions of dark culture were maintained, then embryoids can continuously proliferate and produce new embryoid plexus. This property was important for the mass reproduction of embryoids.

#### Root induction and plant regeneration

The normally developed *D. versipellis* embryoids with cotyledons were transferred to MS+0.5 mg/L IBA+ 0.5 mg/L GA3 medium for the light and dark alternating culture (12 h light/12 h dark). After 2 weeks of culture, the white embryoids turned to green, and further culture can differentiate into complete plants with roots and leaves (Fig. 2 c, o). The plant regeneration rate of the embryoids was >90%.

## Discussion

The growth hormone 2,4-D is an important growth material in plant callus induction and embryoid formation [11, 12], which can induce callus formation, maintain the proliferation of dispersed cells, and contribute to the production of granular callus [13]. In this study, a subculture system of *D. versipellis* callus was also established on a MS medium containing 1 mg/L 2,4-D, and a large number of granular calli was obtained. However, high growth hormone concentrations inhibit the development of granular callus to embryoids [14, 15]. The study on *Dysosma pleiantha* showed that 2,4-D can induce callus production in seeds, but high concentration inhibits callus embryogenesis [16]. When the 2,4-D concentration decreases to 0.1–0.5 mg/L, it has a positive regulatory effect on the induction and proliferation of callus embryoids. The present study also showed similar results.

Cytokinin plays an important role in the growth and development of callus embryoids [17]. Wang et al. showed that the combination of 0.2 mg/L 6-BA and 0.05 mg/L NAA can significantly increase the incidence of callus embryogenesis in *Tapiscia sinensis* Oliv [11]. Another study has also shown that low cytokinin concentrations combined with auxin NAA can realize the subculture and embryoid proliferation under long-term dark culture conditions without calli [18]. In the present study, we used 0.5 mg/L cytokinin 6-BA and 0.1 mg/L auxin NAA to perform embryoid induction and proliferation experiments on the granular callus of *D. versipellis*, which obtained good results, with the embryoid induction rate of >70%. We speculated that the combination of cytokinin 6-BA and auxin NAA may relieve 2,4-D inhibition in the callus embryoid formation stage and promote embryoid development. The results of Kiranmai et al. also suggested that auxin NAA is the most effective hormone in inducing the occurrence of high-frequency embryoids in *Caralluma pauciflora* [19].

Numerous studies have shown that the occurrence of plant embryoids is affected by factors such as genotypes, explants, hormone types and concentrations, metal ions, and other nutrients [9, 20, 21]. Plant callus can be induced to induce effective embryonic development only by determining the suitable culture conditions for plant embryoids. However, this is not a completely 8 controllable growth mode at present. The results of the present study also confirmed this finding. Therefore, in the process of *D. versipellis* tissue culture, the establishment of an efficient, stable, high-frequency embryoid generating system requires further systematic research.

In conclusion, this study systematically studied the callus culture system of *D. versipellis* under specific culture conditions and induced the formation of embryoids and complete plants. This study lays the foundation for further research on the mechanism of action of fungi in promoting the accumulation of podophyllotoxin, which is the main active constituent of the endangered plant *D. versipellis*, as well as the protection of endangered plants and resources and the production of artificial seeds.

## Author Contributions

X.-M. T. and Y.-Q. Z. conceived and designed the experiments. X.-M. T., P.-B. W. and Y. W. performed the experiments. X.-M.T., S.-L. Y. and H.-Z. T. analyzed the data. X.-M. T. and Y.-Q. Z. wrote the manuscript. All authors read and approved the final manuscript.

